# Resolving kinesin stepping: one head at a time

**DOI:** 10.1101/651281

**Authors:** Willi L. Stepp, Zeynep Ökten

**Affiliations:** Physik Department E22, Technische Universität München, James-Franck-Strasse 1, D-85748 Garching, Germany; Munich Center for Integrated Protein Science, D-81377 Munich, Germany

**Keywords:** heteromeric kinesin-2, FIONA, autoinhibition

## Abstract

Kinesins are well-known to power diverse long-range transport processes in virtually all eukaryotic cells. The ATP-dependent processive stepping as well as the regulation of kinesin’ activity have thus been focus of extensive studies over the past decades. It is widely accepted that kinesin motors can self-regulate their activity by suppressing the catalytic activity of the ‘heads’. The distal random coil at the C-terminus, termed ‘tail domain’, is proposed to mediate this autoinhibition, however, a direct regulatory influence of the tail on the processive stepping of kinesin proved difficult to capture. Here, we simultaneously tracked the two distinct head domains in the kinesin-2 motor using dual-color super resolution microscopy (dcFIONA) and reveal for the first time their individual properties during processive stepping. We show that the autoinhibitory wild type conformation selectively impacts one head in the heterodimer but not the other. Our results provide key insights into the regulated kinesin stepping that had escaped experimental scrutiny.

## Introduction

Maintenance of a eukaryotic cell is a daunting task of logistics. One key organizer of the eukaryotic cytoplasm is kinesin, a microtubule-associated molecular motor that transports cargo in diverse settings throughout the cell [1–6]. After association with the trail, kinesin takes many steps in a hand-over-hand fashion with its two head domains and covers micrometer distances in vitro (Figure 1A, top panel). To this end, the motor is propelled by the energy provided by two alternating ATP hydrolysis cycles in the so-called ‘head’ domains [7,8]. Communication between the respective cycles ensures that at least one head remains bound to the microtubule to prevent premature dissociation of the motor from its track [9,10]. The timing of these cycles is characterized by the so-called dwell times, e.g. the time one head remains bound to the filament between steps [11].

**Figure 1.**
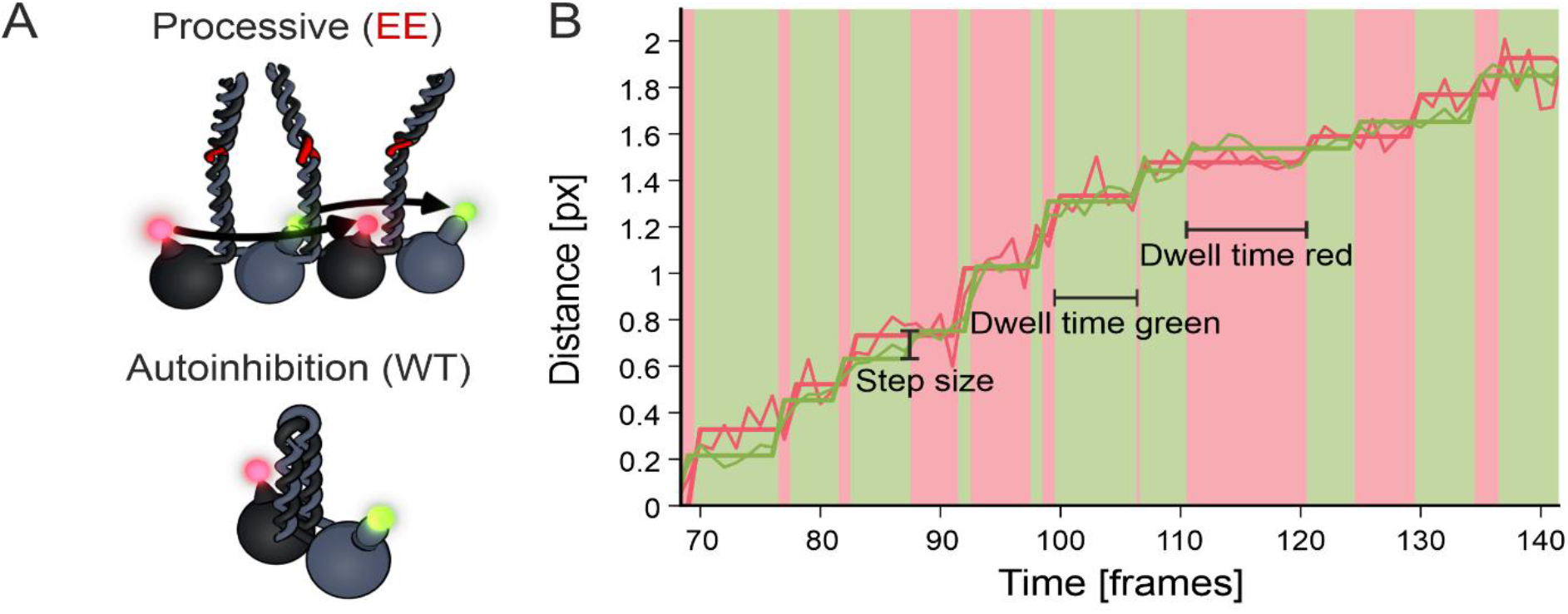
Dual color FIONA setup allows concurrent step detection of both heads in kinesin-2. **A** (top) Depiction of the presumed asymmetric hand-over-hand stepping mode with a heterodimeric kinesin-2 that is labeled with two different fluorophores on its respective head domains. (bottom) Illustration of autoinhibition with the C-terminal tail folded back onto the head domains that in turn suppresses the ATPase activity of the motor. **B** Overlaid stepping of the eeKLP11^Halo^ and the eeKLP20^SNAP^ head domains are shown in green and red, respectively. Data was collected during stepping on microtubules at 0.4 μM ATP. Alternating movement of the motor domains can be seen with corresponding, color coded dwell times highlighted in the background.

Notably, the ATPase activity of kinesin motors can be suppressed by a self-regulatory mechanism termed autoinhibition [12,13]. This is thought to be achieved by folding of the distal C-terminal tail domain onto the N-terminal head domains (Figure 1A, bottom panel). Either removal of the distal tail or preventing the inhibitory folding relieves autoinhibition in vitro (Figure 1A, top panel) [12,14]. Indeed, autoinhibition is proposed to interfere with the entry into a ‘run’ as well as with the stepping of the motor [8–10,12–16]. Competitive binding to cargo or phosphorylation are thought to disengage the tails from the heads in vivo and in vitro [15–17]. Importantly, ectopic activation has been shown to considerably hamper kinesin function in vivo suggesting that self-inhibition is integral to kinesin-dependent transport processes [18]. How the tail-mediated inhibition interferes with the dynamic stepping of a kinesin motor at the molecular level remains an open question [17,19,20].

Several kinesins that belong to the kinesin-2 family form heterotrimeric complexes comprising two distinct motor subunits and one non-motor subunit [15,21–23]. At the example of the heterotrimeric KLP11/20/KAP motor from C. elegans, we previously unmasked the distinct contributions of the KLP11 and KLP20 subunits to the motility and autoinhibition of the heterodimeric KLP11/20 motor in vitro [24]. Indeed, the presence of two kinetically distinct head domains in a kinesin motor long provoked the question of ‘limping’ during the stepping cycles, i.e. a difference in the stepping behavior of the two heads [17,19,25–31]. Limping in kinesin-1 could be enforced under load, however, limping has so far not been resolved with full-length wild type motors during unperturbed stepping [32]. Resolving limping, in particular, necessitates the separation of the dwell time information for each individual head domain within the dimeric motor.

Here, we utilized the heterodimeric nature of the KLP11/20 motor to extract the dwell times from each distinct head simultaneously. To this end, we implemented dual color fluorescence imaging with one nanometer accuracy (dcFIONA) that exposed for the first time the respective dwell times of individual head domains during stepping. The capability to extract information simultaneously from both heads ultimately confirmed the previously suggested limping behavior as well as the inhibitory impact of the tail domain on the stepping of the kinesin motor.

## Results and Discussion

### Dual-color step detection with differentially labeled kinesin-2

To follow the two head domains independently, we introduced SNAP- and Halo-tags at the N-termini of the KLP11/20 heterodimer with wild type stalk (wtKLP11/20 hereafter) and the construct that contained activating mutations in the stalk (KLP11G451E; S452E/ KLP20G444E, G445E; eeKLP11/20 hereafter), respectively (Figure 1A). The corresponding fluorescent JaneliaFluor® dyes of the SNAP- and Halo-tags (JF646 color coded red, JF549 color coded green) labeled the KLP11 and KLP20 subunits with exclusive specificity [33,34] (Suppl. Figure 1).

Using our custom-built setup [21], that we now extended with an additional channel (see Materials and Methods), we performed dcFIONA experiments to track both heads at the same time with exact temporal relation and nanometer resolution. At limiting ATP concentrations, we resolved the stepping of each head individually (Figure 1B). As expected, the step size of the dual-labeled eeKLP11^Halo^/20^SNAP^ was consistent with our previous findings with the eeKLP11^Halo^/20 motor that was labeled on one head domain only (13.2 and13.9 nm vs 13.4 nm [21]) (Suppl. Figure 3).

### The two heads of the KLP11/20 motor display distinct stepping behaviors

The dwell times for kinesin constructs that were labeled on one head only were shown to be distributed according to a double exponential distribution [7,21]. This reflects the ATP-waiting time for the labeled head and the hidden step for the second motor domain [7,21]. In our measurements, we can now extract the dwell times in the ‘step primed’ position, i.e. only the time a head spends in the trailing position before it takes the step.

At limiting ATP concentrations, we measured the individual dwell times of the two heads in the eeKLP11^Halo^/20^Snap^ motor (Figure 1B). For these dual color measurements, one single exponential distribution would be expected for each head [35]. Intriguingly however, we observed two different distributions (Figure 2, left vs right panels). While the dwell times obtained from the KLP20 head domain displayed a single exponential distribution as expected, the dwell times extracted from the KLP11 head domain clearly deviated from a single exponential but were instead consistent with a double exponential distribution (Figure 2, right panels).

**Figure 2.**
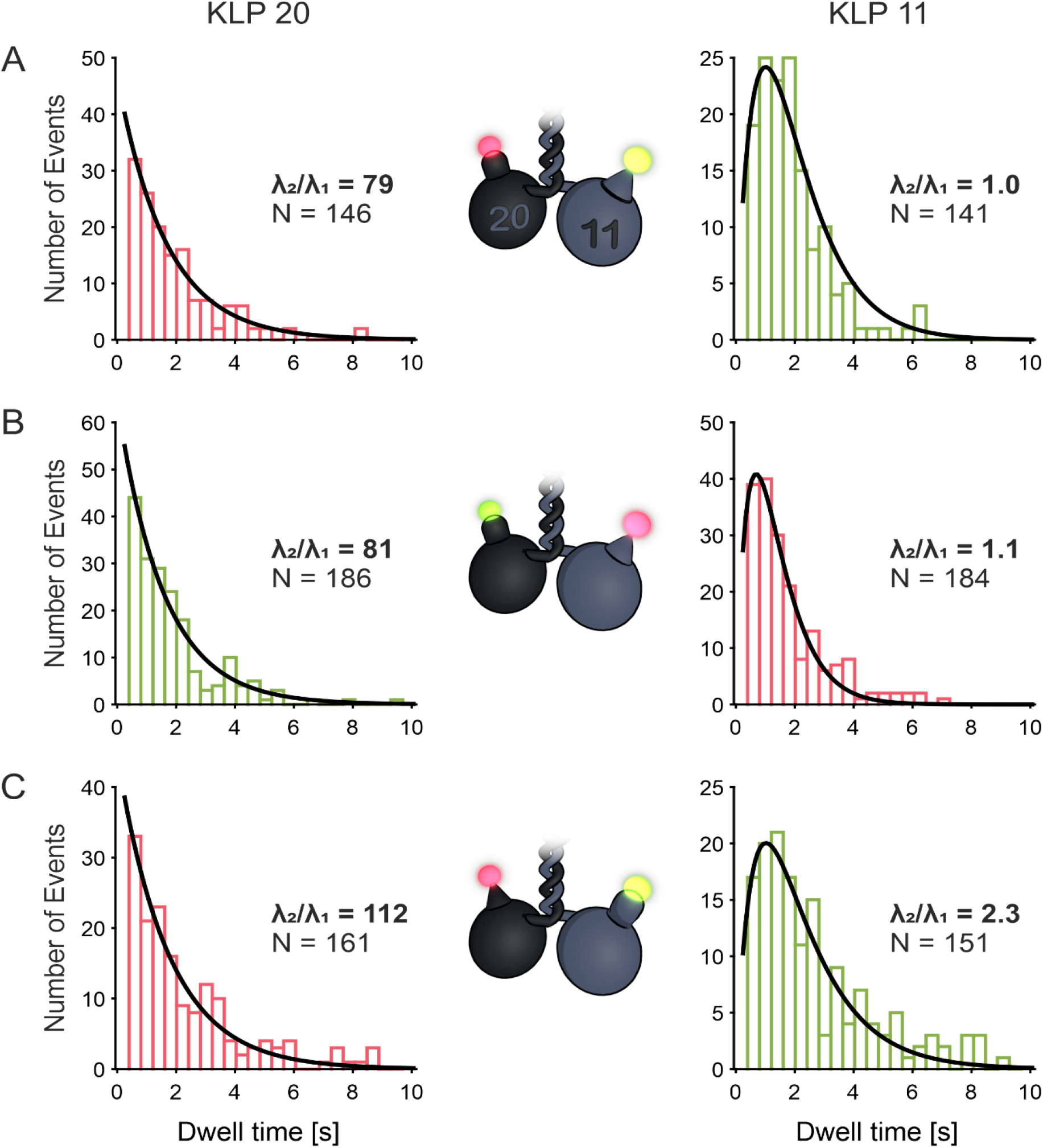
The dwell time distributions of the KLP11 and KLP20 head domains are different. **A-C** Dwell time distributions of KLP20 and KLP11 are depicted with bars in the respective colors (green for JF549, red for JF646) the data was collected in. The width of the bins represents the cycle-time (405 ms), therefore the fits are expected to be independent of the binning. Fits performed with a double exponential model with same settings and starting point (see Supplementary Information for details). KLP20 dwell times show a distribution close to a single exponential distribution (left panels). KLP11 dwell times are fitted well by the double exponential model with similar values for both parameters λ_1_ and λ_2_ (right panels). Fitting of the KLP20 data shown in the left panels with the same model yields a ratio of the two parameters that is about two orders of magnitude higher, resulting in a quasi-single exponential fit. All fits resulted in r-squared values > 90%. **A** eeKLP11Halo/20SNAP (20: λ_1_ = 0.6, λ_2_ = 47.2; 11: λ_1_ = λ_2_ = 1.0) **B** eeKLP11Halo/20SNAP (20: λ_1_ = 0.6, λ_2_ = 48.5; 11: λ_1_ = 1.4, λ_2_ = 1.5) **C** wtKLP11SNAP/20Halo (20: λ_1_ = 0.6, λ_2_ = 67.2; 11: λ_1_ = 0.6, λ_2_ = 1.4) Fitting function: 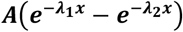

To exclude any influence of the respective fluorophores or their recording by our setup, we switched the dyes (eeKLP11^Halo^/20^SNAP^ vs eeKLP11^Halo^/20^SNAP^) on the respective head domains (Figure 2A vs 2B). In addition, we also switched the position of the Halo- and SNAP-tags themselves (eeKLP11^Halo^/20^SNAP^ vs wtKLP11^SNAP^/20^Halo^) to exclude any influence of the specific tags on the behavior of the motor *per se* (Figure 2B vs 2C). In both cases, we confirmed the double exponential dwell time distribution for the KLP11 head domain (Figure 2, right panels), while the KLP20 head domain consistently displayed a single exponential distribution (Figure 2, left panels).

To further test the consistency of this observation, we fitted both data sets using the same double exponential model (see Supplementary Information). For the KLP11 head domain, the ratio of the two involved parameters was close to 1 (Figure 2, right panels), indicating a similar influence of both values on the stepping behavior. For the KLP20 head domain, in contrast, the ratio was about 100-fold higher, ultimately resulting in a near single exponential fit (Figure 2, left panels).

Together, these findings suggest that the steps taken by the KLP11 head domain include a second rate-limiting event in addition to the waiting time for ATP binding [7]. The observed differences in the dwell time distributions as displayed by the KLP11 and KLP20 head domains ultimately confirm the presumed limping for heterodimeric motors [17,19,25–31]. This behavior of a single head domain could so far not be resolved by tracking the net movement of the motor due to the similar mean dwell times of the two heads [25].

What is the origin of the second rate constant that is displayed specifically by the KLP11 head domain? Notably, our previous work with the wtKLP11/20 suggested an asymmetric autoinhibition mechanism [24]. It required both the presence of the tail and the correct positioning of the KLP11 head within the wtKLP11/20 heterodimer. Strikingly however, solely swapping the positions of the KLP11 and KLP20 head domains sufficed to activate the autoinhibited wtKLP11/20 motor in single molecule and bulk ATPase assays [24,36]. These findings provoke the question whether the presence of the C-termini in the eeKLP11/20 *per se* influences the dwell time distribution of the KLP11 head domain. If true, the autoinhibitory folding in the wtKLP11/20 stalk would be expected to enhance this effect specifically in the KLP11 data (Figure 1A, bottom panel).

### Difference in stepping gives insight into the autoinhibition of kinesin-2

To test this hypothesis, we extracted long dwell times (>2 s) from the KLP11 distributions from Figure 2 (right panels) and refitted them with a single exponential model (Figure 3A). For the wtKLP11/20 motor, the resulting dwell time parameter increased 1.6-fold when compared to the eeKLP11/20 that contains the ATPase activating mutations in the stalk (Figure 3, left panels). This 1.6-fold difference is in fact consistent with the decreased speed of the wild type motor at saturating ATP concentrations (Figure 3B) [20].

**Figure 3.**
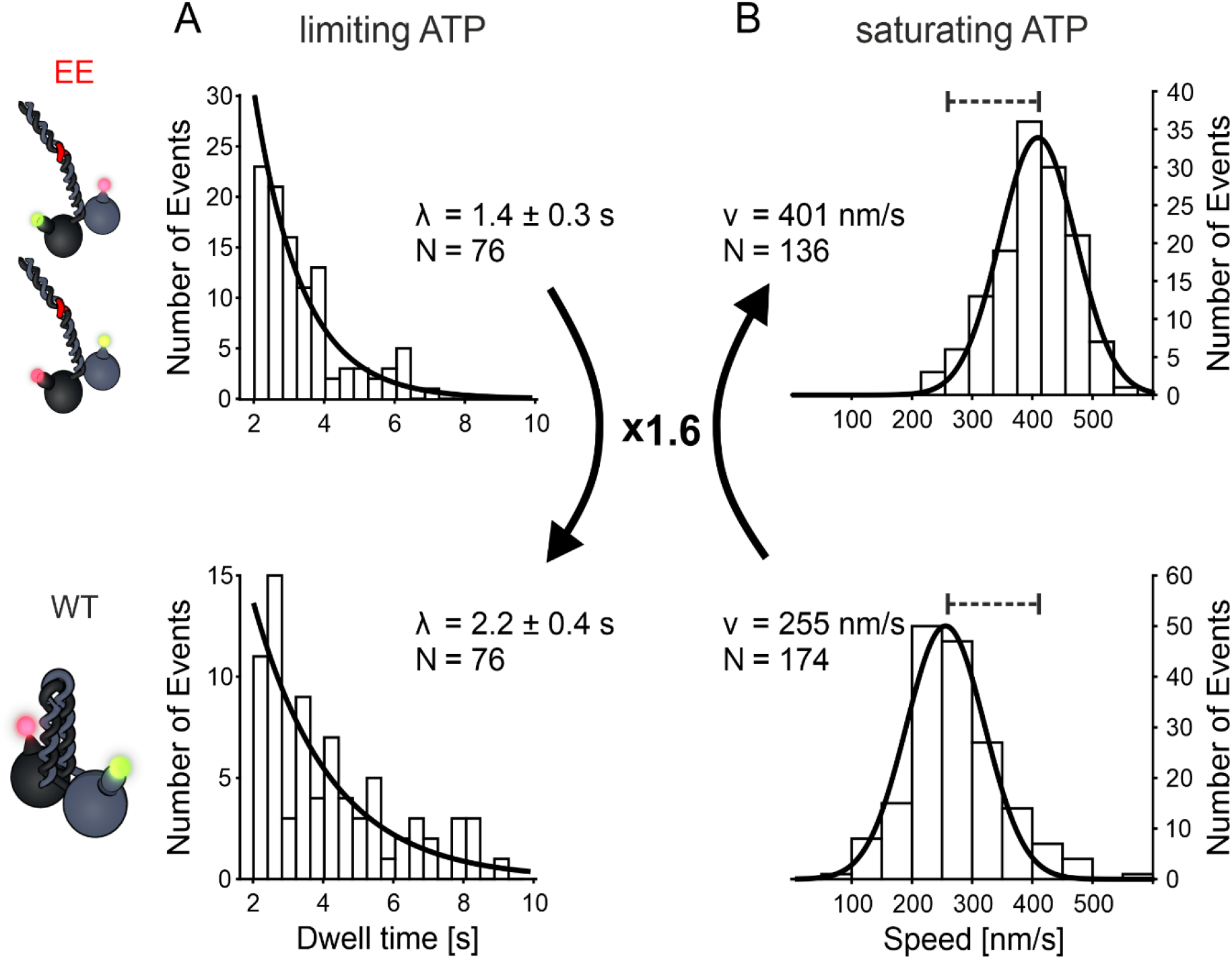
Presence of the wild type C-termini selectively prolongs the dwell times in the stepping of KLP11head domain. **A** Truncated dwell times over 2 s of KLP11 fitted with a single exponential model (data from Figure 2 right panels, A+B eeKLP, C wtKLP). The ratio of the dwell times from the wtKLP11/20 (bottom, 2.2 s) to the eeKLP11/20 (top, 1.4 s) is 1.6. This factor of 1.6 also matches the ratio of mean dwell times as seen in Figure 2 (right panel, A+B eeKLP, C wtKLP). **B** Comparison of velocities from eeKLP11/20 (top, from [21], μ = 90nm/s) and the wtKLP11/20 (bottom, μ = 64 nm/s) at saturating ATP concentrations. The ratio of speeds (eeKLP11/20: 401 nm/s, wtKLP11/20 255 nm/s) is the reversed value of the ratio of dwell times (See Supplementary Figure 2 for run length data of the respective motors).

The first rate constant present in both dwell time distributions is attributed to the ATP-waiting time at low ATP concentrations [7]. Based on previous data [37–39], we speculate that the second rate constant in the dwell time distribution of the KLP11 head domain results from the tail-suppressed ADP release. This effect is strong in the wild type motor in which the flexible hinge in the stalk enables autoinhibitory folding and consequently enhances the ‘head-tail’ interaction (Figure 1A, bottom panel). When the stalk is mutated to prevent autoinhibitory folding (Figure 1A, top panel), the head-tail interaction is hampered, thus selectively shortening the dwell times in the KLP11 stepping (Figure 2, right panels A+B vs. C). Taken together, our capability to distinguish the two head domains during processive stepping provides compelling support for an asymmetric autoinhibition mechanism in the KLP11/20 heterodimer [24]. Indeed, our results unmask for the first time the influence of the tail on the dynamic stepping of a physiological kinesin motor.

Previous tracking of one head domain in the homodimeric kinesin-1 at nanometer resolution using FIONA already represented a major breakthrough given the small 8 nm net displacement of the motor [7]. Being able to trace two head domains simultaneously with the dual color FIONA introduced here now allows the dissection of the *specific* contributions of the head domains to the processive stepping and the regulation thereof at the single molecule level. The next major experimental challenge towards a comprehensive understanding of the kinesin stepping mechanism will be the correlation of the stepping behavior to specific events in the respective ATPase cycles of the motor domains.

## Materials and Methods

### Constructs and design

All constructs were based on the heteromeric kinesin-2 KLP11/20 active in the intraflagellar transport in *C. elegans*. eeKLP mutations were performed as described previously [26]. Halo- and SNAP-tags were fused to the n-terminus of the respective sequences where applicable. The constructs used are:

- eeKLP11^Halo^
- wtKLP11^SNAP^
- eeKLP20^SNAP^
- wtKLP20^Halo^

### Protein expression, purification and fluorescent labeling

All proteins were expressed and purified as described previously [21]. For fluorescent labeling, JaneliaFluor® dyes JF549 and JF646 in Halo- and SNAP-conjugated variants were used [33]. The dyes were mixed 1:1 before incubation and the incubation time with the dyes was prolonged to 90 minutes.

### Microscope setup

Single molecule experiments were performed on a custom-built setup described previously [21]. A 555 nm laser (Oxxius, France) was added to the setup as well as a color split/recombine setup using a high- and a low-pass dichroic to offset the channels on the camera chip.

### Single molecule experiments

Speeds and run length were measured at an ATP concentration of 2 mM. Movies were recorded with an exposure time of 200 ms and 500 frames were recorded before changing the position in the sample.

For step detection experiments, the ATP concentration was reduced to 0.4 μM, the creatin phosphate/creatin phospho kinase system guaranteed stable ATP concentrations over the duration of data collection. Movies were recorded with an exposure time of 400 ms for dual color experiments, resulting in a cycle time of 405 ms.

### Data analysis

All data analysis was performed using ImageJ and custom routines implemented in Matlab (Mathworks Inc.). Traces for speed and run length measurements were extracted by identifying and following peaks depending on their brightness. A position with subpixel accuracy for these traces was assigned using a radial center approach [40]. Runs over several frames were connected by following peaks according to their distance to a peak in the previous frame. Overall distances were calculated with respect to the first detected position in a run. Speeds where then calculated by performing a linear regression on the distance over time data and extracting sequences, that fitted with an r-squared value higher than 95%. Run lengths were determined from the maximum distance from the starting point for each run.

For step detection experiments, a least-squares fit procedure was used to fit a Gaussian profile to the peak data with a starting point deduced from the initial detection of the brightest pixel. This fit provided a higher accuracy subpixel position for each frame, compared to the radial center approach. Due to the lower speeds, the distance over time traces show distinct relocation events. An implementation of the Potts algorithm was used for step detection [41]. Single position spikes in the distance traces were filtered out. The individual sizes of steps were calculated from the mean distances before and after each step.

An algorithm was used to extract sequences of alternating steps in both channels. Dwell times were then calculated from the time of a step in one color to the next step in the other color.

## Acknowledgements

We thank Jonathan B. Grimm and Luke D. Lavis (Janelia Research Campus, HHMI) for their generous gift of the Janelia Fluor compounds. We thank William O. Hancock and Matthias Rief for the helpful comments on the manuscript. The research leading to these results has received funding from the European Research Council (GA no. 335623) to Willi Stepp & Zeynep Ökten and Deutsche Forschungsgemeinschaft SFB863 to Zeynep Ökten.

## Author contributions

WLS, and ZÖ planned the experiments. WLS built the custom TIRF setup and wrote all the customized MATLAB routines. WLS collected and analyzed data. WLS and ZÖ wrote the manuscript.

## Supplementary Information

### Dual exponential fit

In our first impression, the two dwell time distributions for KLP11 and KLP20 were distributed according to a double and single exponential model respectively. To support this assumption, we fitted both distributions with a double exponential model and show, that one of the parameters vanishes for the KLP20 dwell times. The model used was:

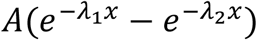

We used the same settings and starting point for the fit:

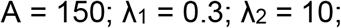

In order to focus on the shorter dwell times, where the effect of the double exponential distribution can be seen best, we introduced a weight function that describes the weight the corresponding data point contributes to the fit:

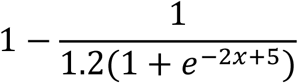

**Supplementary Figure 1.**
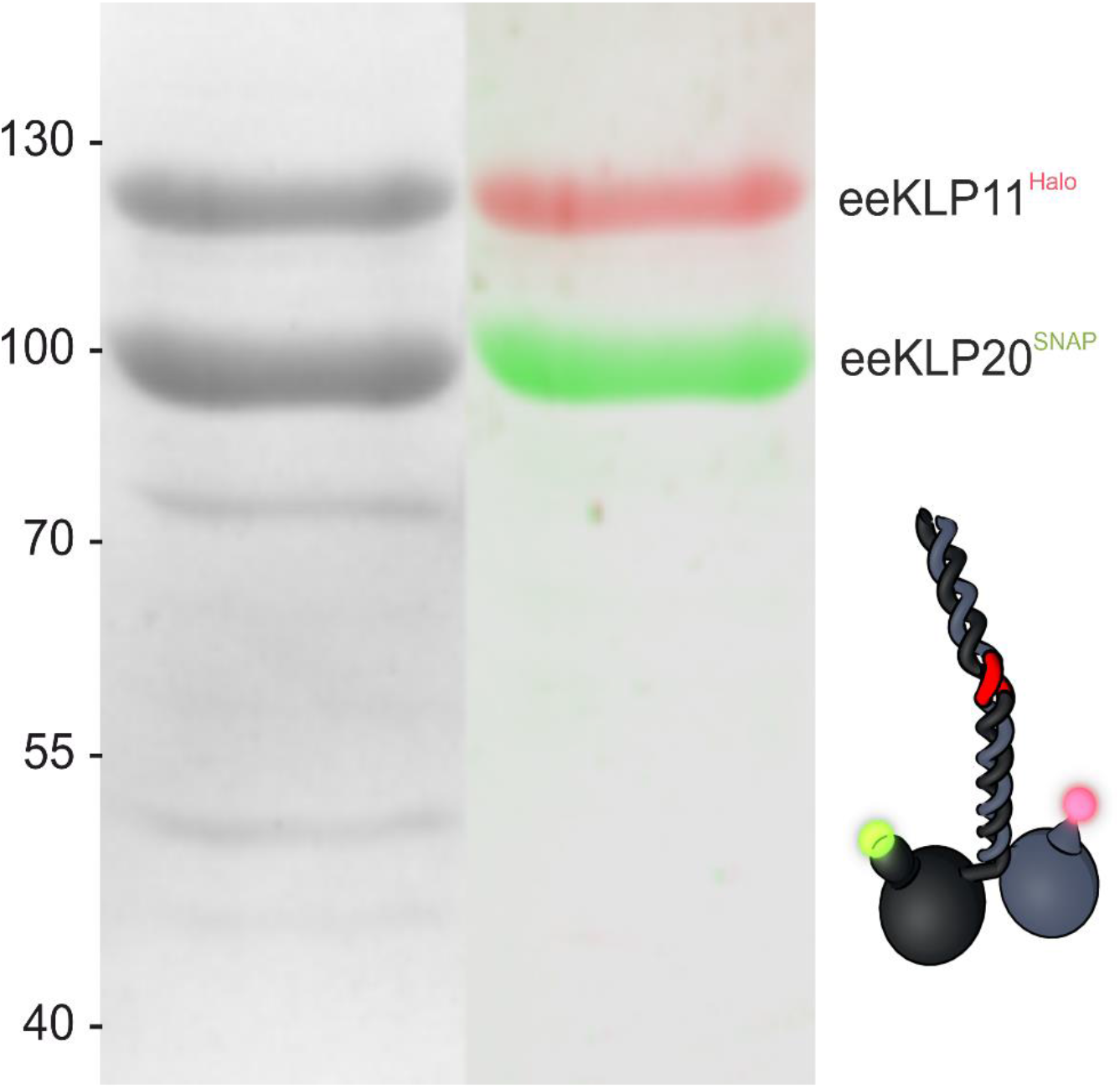
SDS-gel of eeKLP11^Halo^/20^SNAP^ purification shows specific labeling of the tags. The Halo- and Snap-tag was labeled with the respective JaneliaFluor dye respectively. (Left) Coomassie stain of the proteins. (Right) Overlay of two images color coded for the respective channel. Taken on Biostep Celvin S in channels with 525 nm (blue) and 625 nm (red) excitation. The dyes are only detected on the targeted motor domain.

**Supplementary Figure 2.**
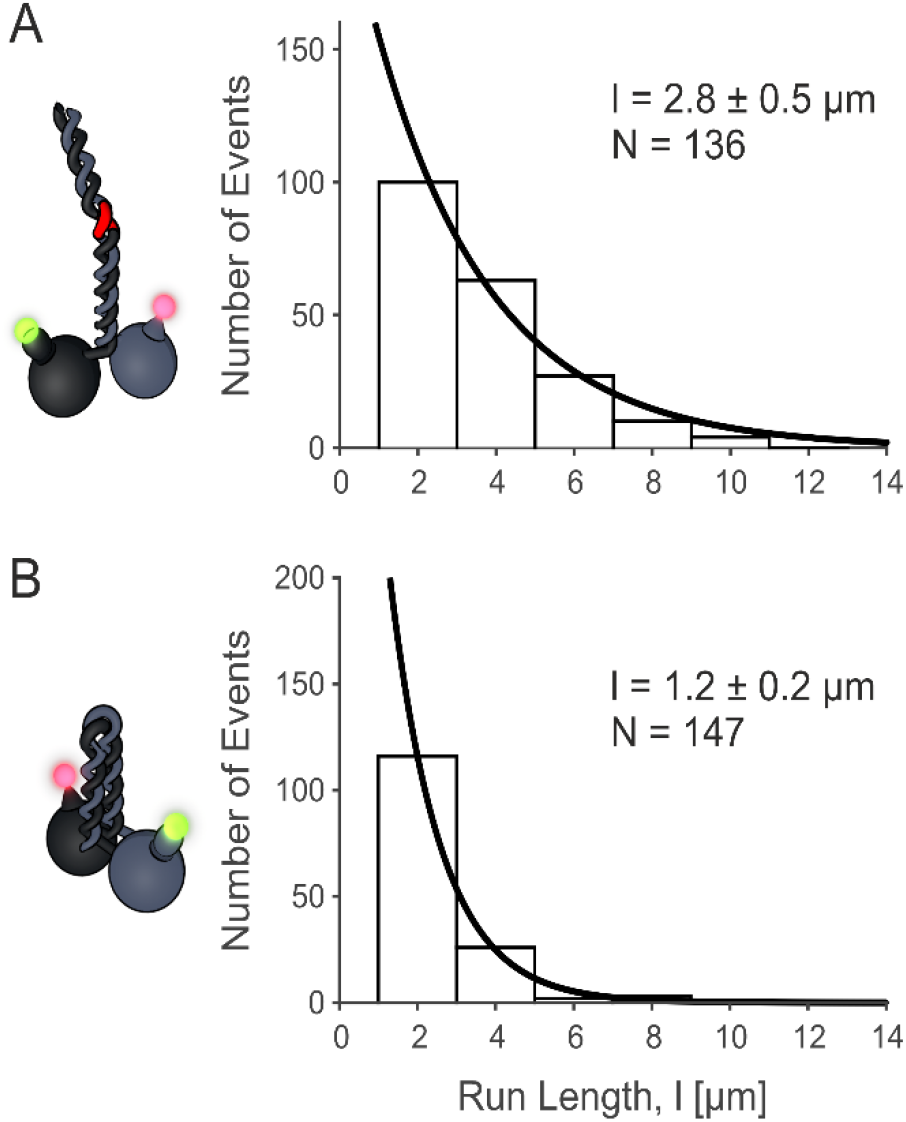
The motors show typical run length. Run length data for (A) eeKLP11^Halo^/20^SNAP^ at 2.8 μm and (B) wtKLP11^SNAP^/20^Halo^ at 1.2 μm. Exponential fit parameter ± 95 % confidence interval.

**Supplementary Figure 3.**
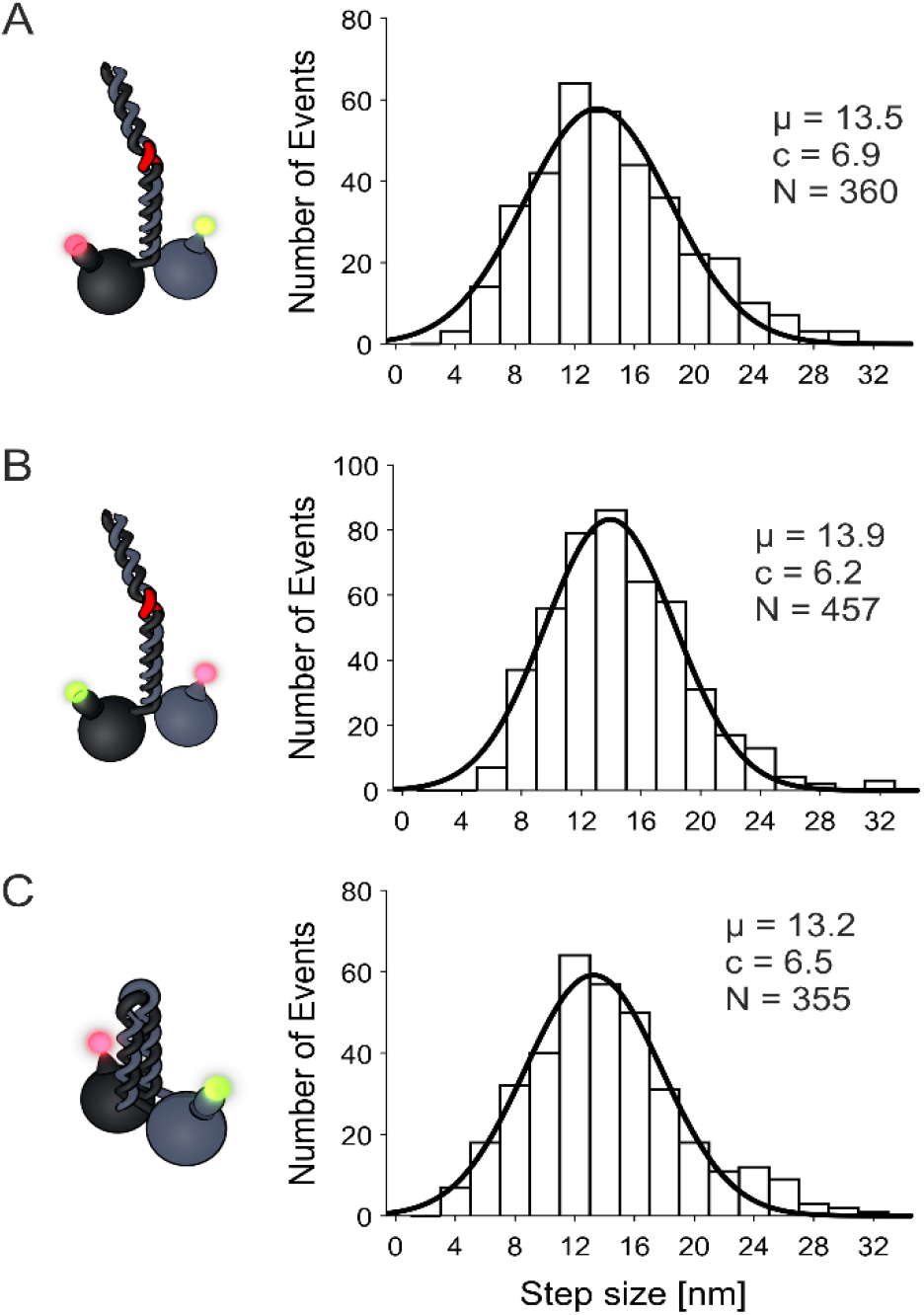
Step sizes of different KLP11/KLP20 constructs are close to what has been measured previously. Step sizes of all runs were combined and pooled from both channels. All results are close to what we measured previously for this motor [21]. **A** eeKLP11^Halo^/20^SNAP^ **B** eeKLP11^Halo^/20^SNAP^ **C** wtKLP11^SNAP^/20^Halo^

